# Perturbed human sub-networks by *Fusobacterium nucleatum* candidate virulence proteins

**DOI:** 10.1101/094136

**Authors:** Andreas Zanzoni, Lionel Spinelli, Shérazade Braham, Christine Brun

**Author notes:** Address correspondence to A. Zanzoni.

## Abstract

*F. nucleatum* is a gram-negative anaerobic species residing in the oral cavity and implicated in several inflammatory processes in the human body. Although *F. nucleatum* abundance is increased in inflammatory bowel disease subjects and is prevalent in colorectal cancer patients, the causal role of the bacterium in gastrointestinal disorders and the mechanistic details of host cell functions subversion are not fully understood.

We devised a computational strategy to identify putative secreted *F. nucleatum* proteins (*Fuso*Secretome) and to infer their interactions with human proteins based on the presence of host molecular mimicry elements. *Fuso*Secretome proteins share similar features with known bacterial virulence factors thereby highlighting their pathogenic potential. We show that they interact with human proteins that participate in infection-related cellular processes and localize in established cellular districts of the host-pathogen interface. Our network-based analysis identified 31 functional modules in the human interactome preferentially targeted by 138 *Fuso*Secretome proteins, among which we selected 26 as main candidate virulence proteins, representing both putative and known virulence proteins. Finally, 6 of the preferentially targeted functional modules are implicated in the onset and progression of inflammatory bowel diseases and colorectal cancer.

Overall, our computational analysis identified candidate virulence proteins potentially involved in the *F. nucleatum* – human cross-talk in the context of gastrointestinal diseases.

## Background

*Fusobacterium nucleatum* is a gram-negative anaerobic bacterium best known as a component of the oral plaque and a key pathogen in gingivitis and periodontitis [1]. It has also been isolated in several inflammatory processes in distinct body sites (*e.g.*, endocarditis, septic arthritis, liver and brain abscesses) and implicated in adverse pregnancy outcomes (reviewed in [2]). Moreover, it has been demonstrated that *F. nucleatum* can adhere to and invade a variety of cell types, thereby inducing a pro-inflammatory response [3–8]. Recent work showed that *(i) F. nucleatum* is prevalent in colorectal cancer (CRC) patients [9–11] and *(ii)* its abundance is increased in new-onset Crohn's disease (CD) subjects [12]. Interestingly, follow-up studies suggested a potential role of this bacterium in CRC tumorigenesis and tumor-immune evasion [13–16].

Despite these findings, a large fraction of *F. nucleatum* gene products are still uncharacterized. Moreover, to date, only a handful of pathogenic factors has been experimentally identified [17,18] and protein interaction data between these factors and human proteins, which could inform on the molecular details underlying host-cell subversion mechanisms, are sparse [4,16,19]. Altogether, this underlines that a comprehensive view of the molecular details of the *F. nucleatum* – human cross-talk is currently missing.

How does *F. nucleatum* could hijack human cells? Pathogens employ a variety of molecular strategies to reach an advantageous niche for survival. One of them consists of subverting host protein interaction networks. Indeed, they secrete and deliver factors such as toxic compounds, small peptides, and even proteins to target the host molecular networks. To achieve this, virulence factors often display structures resembling host components in form and function [20–22] to interact with host proteins, thus providing a benefit to the pathogen [23]. Such “molecular mimics” (*e.g.*, targeting motifs, enzymatic activities and protein-protein interaction elements) allow pathogens to enter the host cell and perturb cell pathways (*e.g.*, [24–26]).

Over the years, several experimental approaches have been applied to identify protein-protein interactions (PPIs) between pathogen and their hosts providing new insights on the pathogen's molecular invasion strategies. However, the vast majority of these systematic studies focused on viruses (e.g., [27–29]) and, to a lesser extent, on bacteria [30–33] and eukaryotic parasites [33,34]. Indeed, as cellular pathogens have large genomes and complex life cycles, the experimental identification of virulence proteins and the large-scale mapping of host-pathogen PPIs require a lot of effort and time [35,36]. In this context, computational approaches have proved to be instrumental for the identification of putative pathogenic proteins (*e.g.*, [37,38]), the characterization of molecular mimics [23,39,40] and the inference of their interactions with host proteins (for a review see [41]).

Here, in order to gain new insights on the molecular crosstalk between *F. nucleatum* and the human host, we devised a computational strategy combining secretion prediction, protein-protein interaction inference and protein interaction network analyses (Figure 1). Doing so, we defined a secretome of the bacterium and the human proteins with which they interact based on the presence of mimicry elements. We identified the host cellular pathways that are likely perturbed by *F. nucleatum* including immune and infection response, homeostasis, cytoskeleton organization and gene expression regulation. Interestingly, our results identify candidate virulence proteins, including the established Fap2 adhesin, and provide new insights underlying the putative causative role of *F. nucleatum* in colorectal cancer and inflammatory bowel diseases.

**Figure 1.**
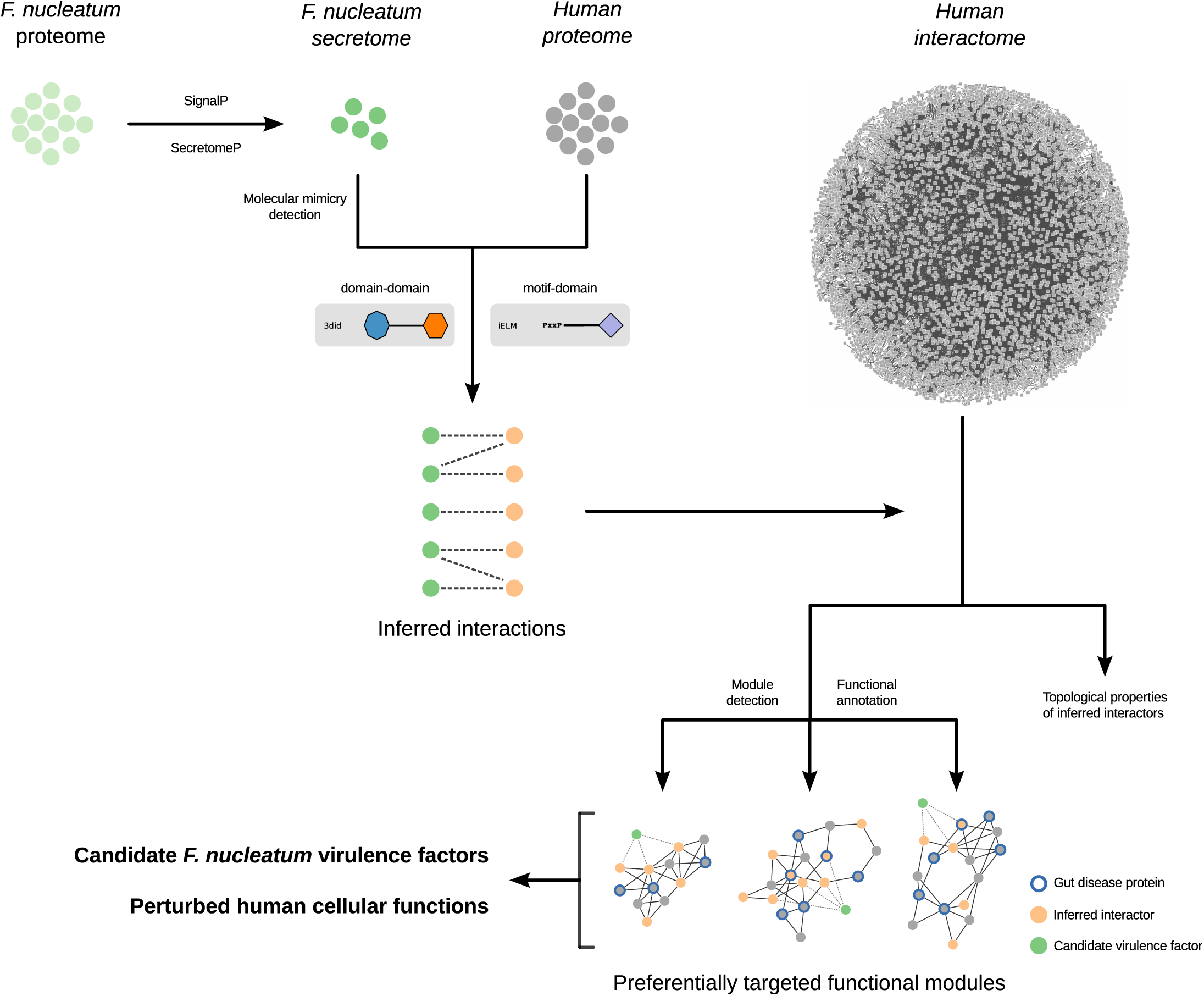
Flow strategy of our computational approach.

## Results

### Prediction of *F. nucleatum* secreted proteome

Previous computational analyses highlighted that *F. nucleatum* has a reduced repertoire of secretion machinery [42,43] meaning that it might exploit alternative “non-classical” translocation mechanisms to unleash virulence proteins. Thus, we sought to identify putative *F. nucleatum* secreted proteins by analyzing the 2046 protein sequences of the type species *F. nucleatum subsp. nucleatum* (strain ATCC 25586) proteome using two distinct algorithms: SignalP [44] for peptide-triggered secretion and SecretomeP [45] for leaderless protein secretion. While the SignalP algorithm predicted 61 *F. nucleatum* sequences being secreted *via* classical/regular secretion pathways, SecretomeP found 176 proteins as possibly secreted through non-classical routes. In total, we identified 237 putative secreted proteins in the *F. nucleatum* proteome (herein called “*Fuso*Secretome”) (see Table S1). Notably, we were able to correctly predict as secreted all the *F. nucleatum* virulence proteins known so far, namely FadA (FN0264), Fap2 (FN1449), RadD (FN1526) and the recently identified Aid1 adhesin (FN1253) [46]. This result underlines the relevance of secretion prediction to identify novel putative virulence proteins in the *F. nucleatum* proteome.

It has been shown that disorder propensity is an emerging hallmark of pathogenicity [47,48]. As SecretomeP exploits protein disorder as a predicting feature, we analyzed the intrinsic disorder content of the *Fuso*Secretome proteins identified by the SignalP algorithm only. We indeed observed a significantly higher disorder propensity of these proteins compared to the non-secreted proteins (P-value = 1.9×10^−4^, Kolgomorov-Smirnov test, two-sided) (Figure 2; Figure S1; Table S2), further reinforcing the possible role of the *Fuso*Secretome in the infection/invasion process.

**Figure 2.**
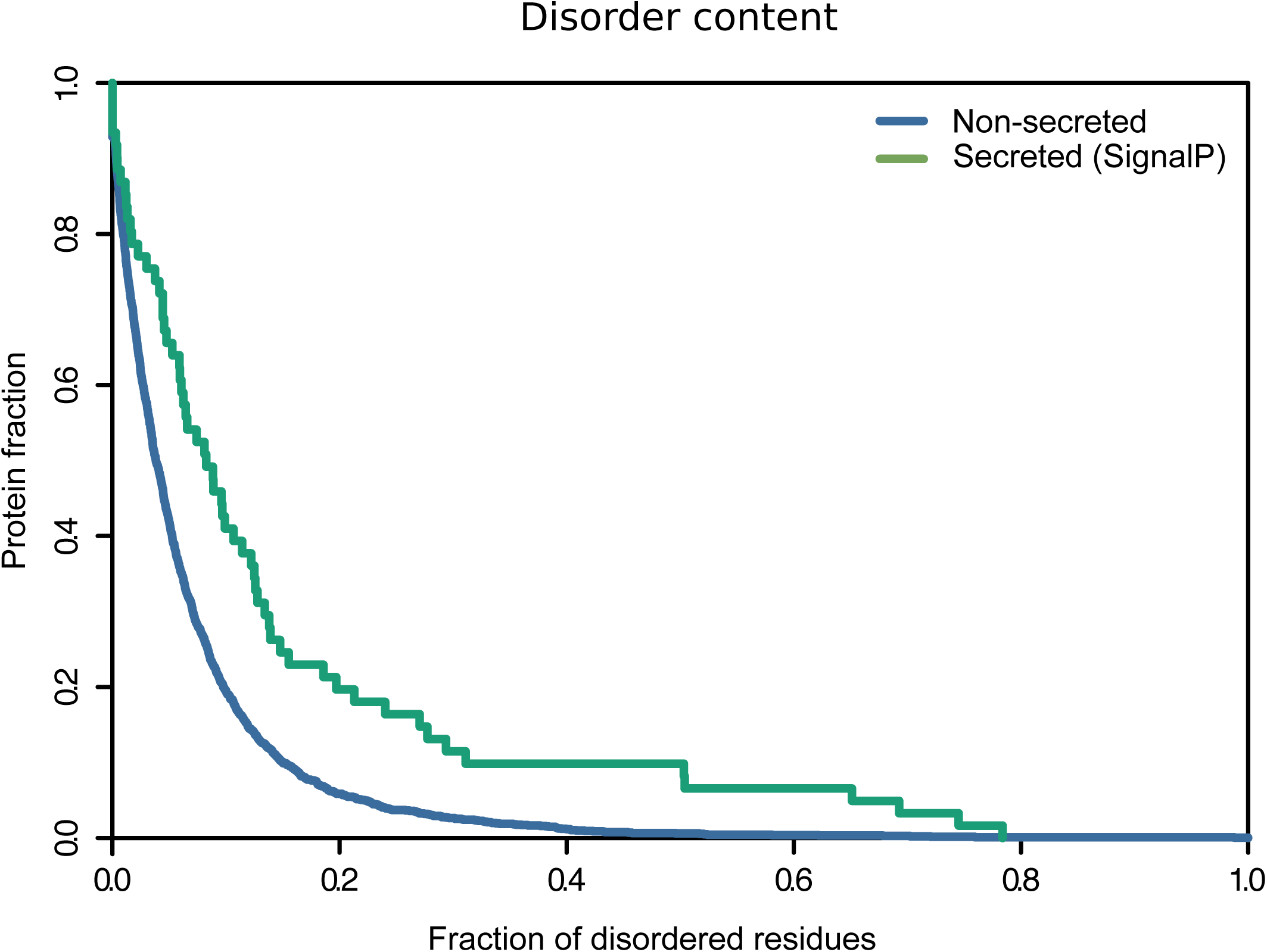
Disorder propensity of the *Fuso*Secretome. SignalP-secreted proteins show a significantly higher fraction of disordered residues compared to non-secreted proteins (P-value = 1.9×10^−4^, Kolgomorov-Smirnov test, two-sided).

To detect functional elements that can further contribute to *F. nucleatum* pathogenicity, we sought for the presence of globular domains in the *Fuso*Secretome. We observed an enrichment of domains mainly belonging to the outer membrane beta-barrel protein superfamily (Table 1). Six out of the eight over-represented domains among the *Fuso*Secretome proteins are also found in known virulence proteins of gram-negative bacteria [49] and are involved in adhesion, secretion, transport and invasion. Altogether, these findings suggest that *Fuso*Secretome proteins display features of known virulence proteins and can likely be involved in the cross-talk with the human host.

**Table 1.** Enrichment of Pfam domains in the *Fuso*Secretome compared to non-secreted proteins.

| Pfam domain | Pfam Clan<sup>a</sup> | Function | VFDB<sup>b</sup> | Human<sup>c</sup> | # *Fuso*Secretome<sup>d</sup> | # non-secreted<sup>e</sup> | Corrected P-value<sup>f</sup> |
| --- | --- | --- | --- | --- | --- | --- | --- |
| MORN repeat variant | MORN repeat | Invasion | - | - | 16 | 2 | 2.17x10<sup>-12</sup> |
| Autotransporter beta-domain | Outer membrane beta-barrel protein superfamily | Adhesion | ✓ | - | 9 | 0 | 2.4x10<sup>-7</sup> |
| Haemolysin secretion/activation protein ShlB/FhaC/HecB | Outer membrane beta-barrel protein superfamily | Secretion | ✓ | - | 5 | 0 | 0.002 |
| TonB-dependent Receptor Plug Domain | Ubiquitin superfamily | Transport | ✓ | - | 5 | 0 | 0.002 |
| TonB dependent receptor | Outer membrane beta-barrel protein superfamily | Transport | ✓ | - | 4 | 0 | 0.011 |
| Surface antigen variable number repeat | POTRA domain superfamily | Folding | - | - | 4 | 0 | 0.011 |
| YadA-like C-terminal region | Pilus subunit | Adhesion | ✓ | - | 4 | 0 | 0.011 |
| Haemagglutinin repeat | Pectate lyase-like beta helix | Adhesion | ✓ | - | 4 | 0 | 0.011 |
| Bacterial extracellular solute-binding proteins, family 5 Middle | Periplasmic binding protein clan | Transport | - | - | 5 | 4 | 0.082 |
| Coiled stalk of trimeric autotransporter adhesion | - | Adhesion | ✓ | - | 3 | 0 | 0.09 |
| Pyruvate flavodoxin/ferredoxin oxidoreductase, thiamine diP-bdg | Thiamin diphosphate-binding superfamily | Metabolism | - | - | 3 | 0 | 0.09 |
| TPR repeat | Tetratricopeptide repeat superfamily | Protein binding | ✓ | ✓ | 6 | 8 | 0.094 |

<sup>a</sup> A clan is defined as a collection of related Pfam entries sharing sequence or structural similarity. <sup>b</sup> Pfam entry detected in at least one protein sequence stored in the database of known bacterial virulence factors. <sup>c</sup> Pfam entry detected in at least one human protein sequence. <sup>d</sup> Number of occurrences in the *Fuso*Secretome. <sup>e</sup> Number of occurrences in non-secreted proteins.
<sup>f</sup> Pfam domain matches with a corrected P-value <0.1.

### Inference of the *Fuso*Secretome – human interaction network

Generally, pathogens employ a variety of molecular strategies to interfere with host-cell networks, controlling key functions such as plasma membrane and cytoskeleton dynamics, immune response and cell death/survival. In particular, their proteins often carry a range of *mimics*, which resemble structures of the host at the molecular level, to ‘sneak’ into host cells [20–22,50].

Here, we focused on putative molecular mimicry events that can mediate the interaction with host proteins: *(i)* globular domains that occur in both *Fuso*Secretome and the human proteome and *(ii)* known eukaryotic short linear motifs (SLiMs) found in *Fuso*Secretome proteins. SLiMs are short stretches of 3-10 contiguous amino acids residues that often mediate transient PPIs and tend to bind with low affinity [51].

We first scanned the sequences of the *Fuso*Secretome and human proteins for the presence of domains as defined by Pfam [52]. We identified 55 “host-like” domains in 50 *Fuso*Secretome proteins out of 237, including several domains related to ribosomal proteins, aminopeptidases and tetratricopeptide repeats (TPR) (Table S3). Interestingly, 29 of these domains are also found in known bacterial binders of human proteins [30].

We next detected the occurrence of experimentally identified SLiMs gathered from the Eukaryotic Linear Motif (ELM) database [53]. As linear motifs are short and degenerate in sequence, SLiM detection is prone to over-prediction [54]. To reduce the number of false positives, we kept occurrences falling in conserved and disordered protein sequences (see Methods). Indeed, known functional SLiMs show a higher degree of conservation compared to surrounding residues [51] and are located in unstructured regions [55,56]. In this way, we identified at least one putative mimicry SLiM in 139 *Fuso*Secretome proteins. Most of the 57 different detected SLiMs represents binding sites such as motifs recognized by PDZ, SH3 and SH2 domains (Table S3).

We exploited these putative mimicry events to infer the interaction with human proteins by using templates of domain-domain and SLiM-domain interactions (see Methods for further details). Doing so, we obtained 3,744 interactions (1,544 domain- and 2,201 SLiM-mediated interactions, respectively) between 144 *Fuso*Secretome, which we designated as “candidate virulence proteins”, and 934 human proteins (Table S4 and Table S5) designated as “human inferred interactors”.

In order to assess the reliability of the inferences, we evaluated the biological relevance of the putative human interactors by performing enrichment analyses of different orthogonal datasets using as a reference background all the proteins encoded by the human genome.

First, human proteins experimentally identified as binders or targets of bacterial and viral proteins are over-represented among the 934 inferred human interactors of the *Fuso*Secretome proteins (415 proteins, 1.3-fold, P-value = 1.61 × 10^−11^). Notably, the over-representation holds when bacterial and viral binders are considered separately (176 bacterial interactors, 1.1-fold, P-value = 3.5×10^−3^ and 338 viral interactors, 1.5-fold, P-value < 2.2×10^−16^). This result is consistent with current knowledge on convergent targeting of host proteins by distinct pathogens [30,33,57,58]. Second, according to the Human Proteins Atlas (see Methods), the vast majority of the inferred human interactors has been detected either in small intestine (652, 70%) or colorectal (671 proteins, 72%) tissues as well as in the saliva (673, 72%), confirming their presence in human body sites hosting *F. nucleatum.* Third, we assessed whether the inferred human interactors are implicated in gastrointestinal disorders by seeking for an over-representation of genes associated to such diseases (see Methods). Indeed, the human interactors of the *Fuso*Secretome are enriched in *(i)* proteins identified in the human colon secretomes of colorectal cancer (CRC) tissue samples (3.5-fold, P-value < 2.2 × 10^−16^), *(ii)* proteins encoded by genes whose expression correlates with *F. nucleatum* abundance in CRC patients [13] (2-fold, P-value = 4×10^−4^) and *(iii)* genes associated with inflammatory bowel diseases (IBDs) (2-fold, P-value = 8×10^−4^). We obtained very similar enrichments by using a reduced statistical background corresponding to the interaction inference space (see Methods and Additional file 12: Supplementary Results).

Altogether, the results of these analyses highlight the relevance of the inferred human interactors as putative binders of *Fuso*Secretome proteins and their potential implication in gut diseases, therefore validating the undertaken inference approach.

### Functional role of the human proteins targeted by *F. nucleatum*

Globally, the inferred *Fuso*Secretome human interactors are involved in several processes related to pathogen infection such as immune response and inflammation, response to stress, endocytosis as shown by the 137 significantly enriched Biological Processes Gene Ontology (GO) terms among their annotations (Table 2, Table S6A). Similarly, the targeted human proteins are over-represented in 125 pathways (54 from KEGG and 71 from Reactome databases [59,60]) involved in cell adhesion and signaling, extracellular matrix remodeling, immunity, response to infection and cancer-related pathways (Table 2, Table S6A). These human proteins are mainly localized in the extracellular space, plasma membrane and at cell-cell junctions that represent the main districts involved in the initial encounter between a pathogen and the host, as indicated by the over-representation of 30 Cellular Component GO terms (Table 2, Table S6A). A substantial fraction of these enriched functional categories is significantly over-represented when using the reduced statistical background as well (see Methods and Table S6B). Overall, this indicates that our inferred interactions can participate in the *F. nucleatum* – human crosstalk.

**Table 2.** Significant Gene Ontology and pathways annotations among FusoSecretome inferred human interactors.

| Annotation source | Annotation ID | Annotation name | Corrected P-value |
| --- | --- | --- | --- |
| *Biological Process* | GO:0006415 | translational termination | 7.57x10<sup>-41</sup> |
|  | GO:0006414 | translational elongation | 2.65x10<sup>-34</sup> |
|  | GO:0006457 | protein folding | 2.51x10<sup>-29</sup> |
|  | GO:0006413 | translational initiation | 6.63x10<sup>-27</sup> |
|  | GO:0072376 | protein activation cascade | 1.17x10<sup>-25</sup> |
|  | GO:0051604 | protein maturation | 5.45x10<sup>-23</sup> |
|  | GO:0006614 | SRP-dependent cotranslational protein targeting to membrane | 1.93x10<sup>-22</sup> |
|  | GO:0006956 | complement activation | 3.62x10<sup>-21</sup> |
|  | GO:0002697 | regulation of immune effector process | 4.03x10<sup>-18</sup> |
|  | GO:0030449 | regulation of complement activation | 4.10x10<sup>-18</sup> |
| *Cellular Component* | GO:0005840 | ribosome | 9.81x10<sup>-37</sup> |
|  | GO:0022626 | cytosolic ribosome | 2.76x10<sup>-23</sup> |
|  | GO:0005912 | adherens junction | 4.15x10<sup>-22</sup> |
|  | GO:0098552 | side of membrane | 2.46x10<sup>-18</sup> |
|  | GO:0030055 | cell-substrate junction | 2.85x10<sup>-17</sup> |
|  | GO:0072562 | blood microparticle | 9.20x10<sup>-17</sup> |
|  | GO:0005761 | mitochondrial ribosome | 4.28x10<sup>-12</sup> |
|  | GO:0019897 | extrinsic component of plasma membrane | 2.35x10<sup>-09</sup> |
|  | GO:0005911 | cell-cell junction | 4.54x10<sup>-08</sup> |
|  | GO:0031012 | extracellular matrix | 3.91x10<sup>-07</sup> |
| *KEGG* | KEGG:03010 | Ribosome | 4.34x10<sup>-39</sup> |
|  | KEGG:04610 | Complement and coagulation cascades | 2.28x10<sup>-22</sup> |
|  | KEGG:04514 | Cell adhesion molecules (CAMs) | 2.88x10<sup>-09</sup> |
|  | KEGG:04141 | Protein processing in endoplasmic reticulum | 2.03x10<sup>-08</sup> |
|  | KEGG:04660 | T cell receptor signaling pathway | 1.33x10<sup>-07</sup> |
|  | KEGG:05150 | Staphylococcus aureus infection | 1.78x10<sup>-06</sup> |
|  | KEGG:04380 | Osteoclast differentiation | 7.25x10<sup>-06</sup> |
|  | KEGG:05203 | Viral carcinogenesis | 8.21x10<sup>-06</sup> |
|  | KEGG:05169 | Epstein-Barr virus infection | 1.11x10<sup>-05</sup> |
|  | KEGG:05164 | Influenza A | 1.77x10<sup>-05</sup> |
| *Reactome* | REAC:1592389 | Activation of Matrix Metalloproteinases | 4.77x10<sup>-28</sup> |
|  | REAC:192823 | Viral mRNA Translation | 2.14x10<sup>-19</sup> |
|  | REAC:156902 | Peptide chain elongation | 2.14x10<sup>-19</sup> |
|  | REAC:975956 | Nonsense Mediated Decay independent of the Exon Junction Complex | 1.83x10<sup>-17</sup> |
|  | REAC:977606 | Regulation of Complement cascade | 5.13x10<sup>-16</sup> |
|  | REAC:202733 | Cell surface interactions at the vascular wall | 7.22x10<sup>-16</sup> |
|  | REAC:5368287 | Mitochondrial translation | 3.61x10<sup>-14</sup> |
|  | REAC:3371453 | Regulation of HSF1-mediated heat shock response | 3.92x10<sup>-14</sup> |
|  | REAC:3371599 | Defective HLCS causes multiple carboxylase deficiency | 1.33x10<sup>-08</sup> |
|  | REAC:420597 | Nectin/Necl trans heterodimerization | 1.33x10<sup>-08</sup> |

For each annotation source, the ten most significant terms are reported. The full list of annotation enrichments is available in Table S6.

### *F. nucleatum* targets topologically important proteins in the host network

To gain a broader picture of the inferred interactions in the cellular context, we mapped the *Fuso*Secretome human interactors on a binary human interactome built by gathering protein interactions data from both small-scale experiments and systematic screens reported in the literature (see Methods and Table S7). Around 70% of the inferred human interactors (*i.e.*, 663 proteins) are present in the human binary interactome. Interestingly, the human targeted proteins occupy topologically important positions in the interactome as shown by their significantly higher number of interactions and higher values of betweenness centrality compared to other network proteins (number of interactions: mean = 23 vs. 11, P-value = 1.9×10^−10^; betweeness centrality: mean = 0.00078 vs. 0.00018, P-value = 6.2×10^−12^; two-sided Mann-Whitney U test) (Figure 3).

**Figure 3.**
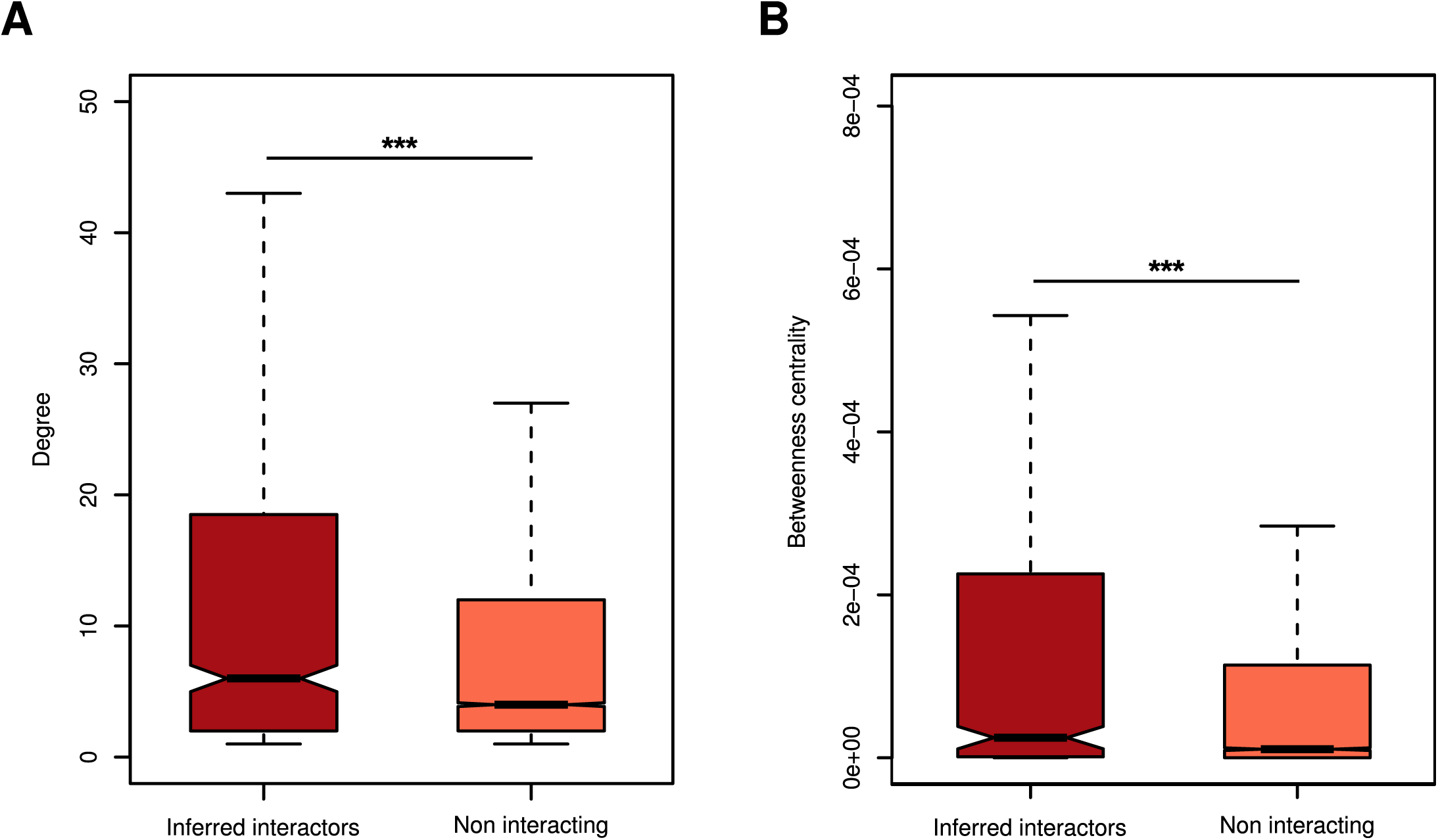
Topological properties of inferred human interactors in the human interactome. (A) Inferred human interactors have more interaction partners and (B) higher values of betwenness centrality compared to non-interacting proteins in the human interactome.

The human interactome is composed of functional network modules, defined as group of proteins densely connected through their interactions and involved in the same biological process [61] (see Methods). We thus next investigated the 855 functional modules that we previously detected [62] using the OCG algorithm that decomposes a network into overlapping modules, based on modularity optimization [63] (Table S8). A significant number of interactors participate in 2 or more of these functional units (259 proteins, 1.3-fold enrichment, P-value = 1.4 × 10^−7^), indicating that the *Fuso*Secretome tends to target multifunctional proteins in the human interactome [63]. Moreover, among the multifunctional inferred human interactors we found an enrichment of extreme multifunctional proteins (52 interactors, 2-fold enrichment, P-value = 1.0×10^−5^), which are defined as proteins involved in unrelated cellular functions and may represent candidate moonlighting proteins [64]. This suggests that the *Fuso*Secretome might perturb multiple cellular pathways simultaneously by targeting preferentially a whole range of multifunctional proteins.

### Functional subnetworks of the human interactome perturbed by *F. nucleatum* and identification of the main candidate virulence proteins

Based on their enrichment in inferred human interactors, 31 network modules (∼4% of the 855 detected modules) are preferentially targeted by 138 distinct proteins of the *Fuso*Secretome (Table 3). Targeted modules are involved in relevant processes such as immune response, cytoskeleton organization, cancer and infection-related pathways (Table 3 and Table S9). Moreover, proteins belonging to these modules are mainly localized in the extracellular space or in membranous structures (Table 3 and Table S9), which represent important districts of the microbe-host interface. Interestingly, the enrichment of functional categories related to gene expression regulation (Table S9) in several modules suggests novel potential host subversion mechanisms by *F. nucleatum*.

**Table 3.**
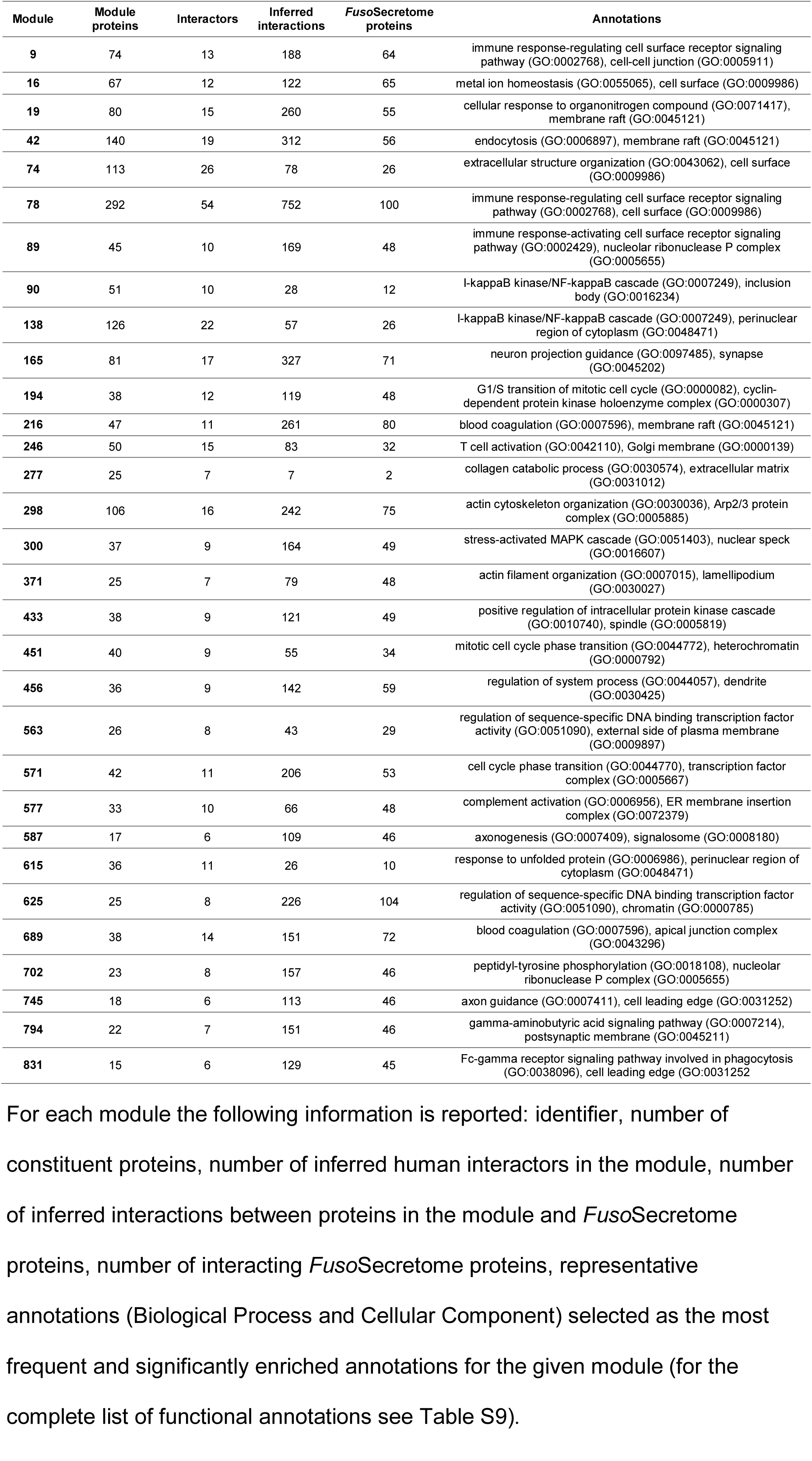
Network module significantly enriched in inferred human interactors.

These modules are targeted on average by 50 FusoSecretome proteins (ranging from 2 to 104 per module) and the number of inferred host-pathogen interactions for each module varies considerably (Table 3). What are the main network perturbators among the *Fuso*Secretome proteins? To quantify their impact on network modules based on the number of interactions they have with each of them, we computed a Z-score (see Methods, Table S10). We considered the 26 *Fuso*Secretome proteins having a perturbation Z-score > 2 in at least one module as main candidate virulence proteins. They consist in outer membrane proteins, enzymes, iron-binding proteins and protein involved in transport (Table 4). Ten of them (38%) can perturb at least two distinct modules (Figure 4A). Notably, we identified among the candidates, the known virulence protein Fap2 (FN1449) (Figure 4B) that targets 4 modules, and a protein containing the MORN_2 domain (FN2118) (Figure 4C) recently identified as a key element in actively invading *F. nucleatum* species [65], which perturbs 6 modules. On the other hand, 25 preferentially targeted modules are perturbed by at least two candidate virulence proteins, Module 78 involved in immune response being the most potentially subverted (Figure 4A).

**Figure 4.**
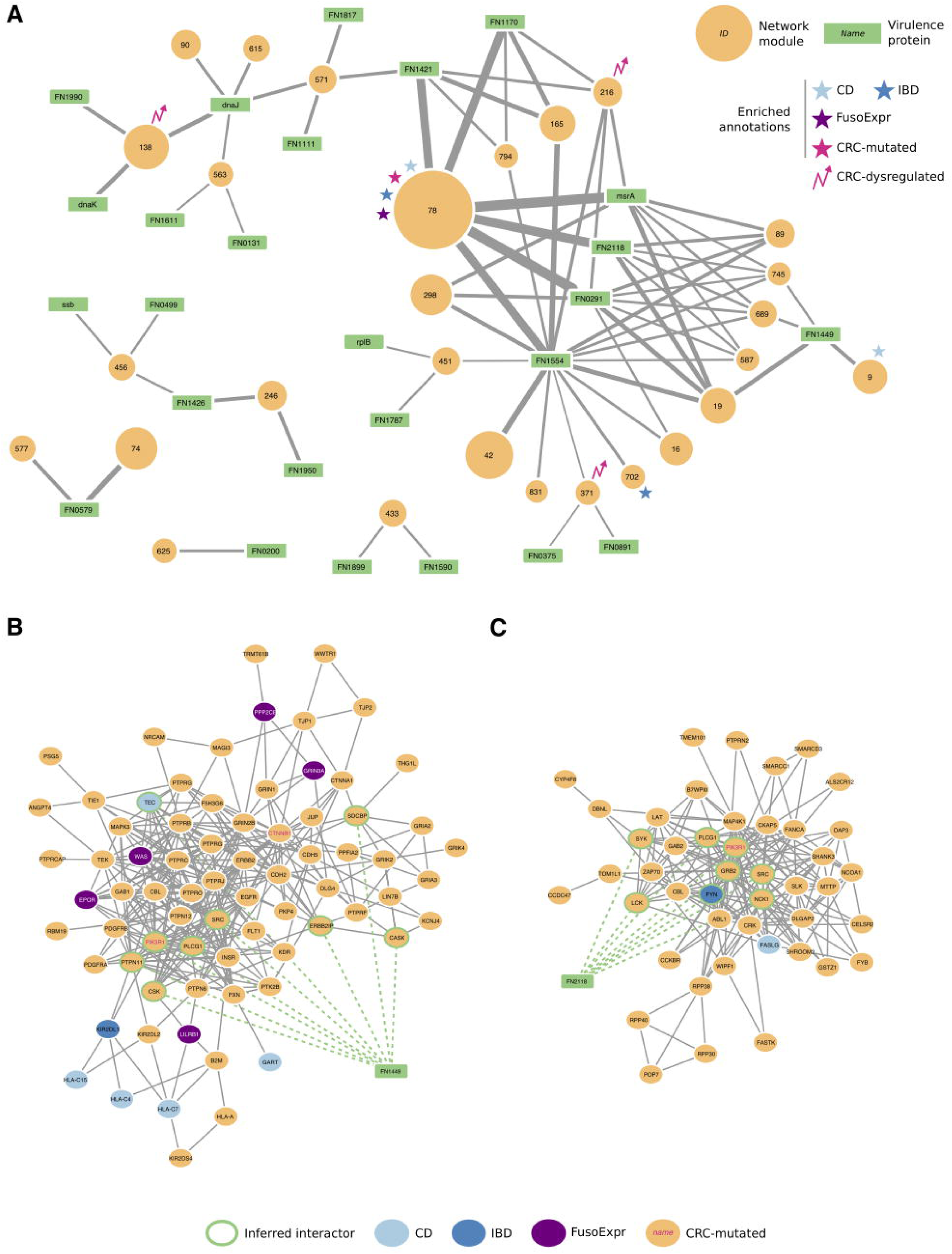
Interaction network between *Fuso*Secretome candidate virulence proteins and preferentially targeted modules. (A) Candidate virulence proteins are depicted as green rectangular nodes labelled with respective gene symbol, whereas network modules as orange circles, whose size is proportional to the number of proteins belonging to each module and are labelled with the corresponding identifier. Edge width is proportional to the number of inferred interactions of a virulence protein with a given module. Network modules enriched in gut-related disease gene sets are labelled with symbols of different colors (i.e., light blue star: Crohn's disease, CD; dark blue star: Inflammatory bowel disease, IBD; violet star: genes whose expression correlates with *F. nucleatum* abundance in colorectal cancer patients, FusoExpr; rose star: genes mutated in colorectal cancer, CRC-mutated; rose zig-zag arrow: dysregulated expression during colorectal cancer progression, CRC-dysregulated). (B) The protein Fap2 (FN1449) interacts with 9 proteins (nodes with a green border) of Module 9 and (C) the MORN2 domain containing protein (FN2118) interacts with 8 proteins in Module 89.

**Table 4.**
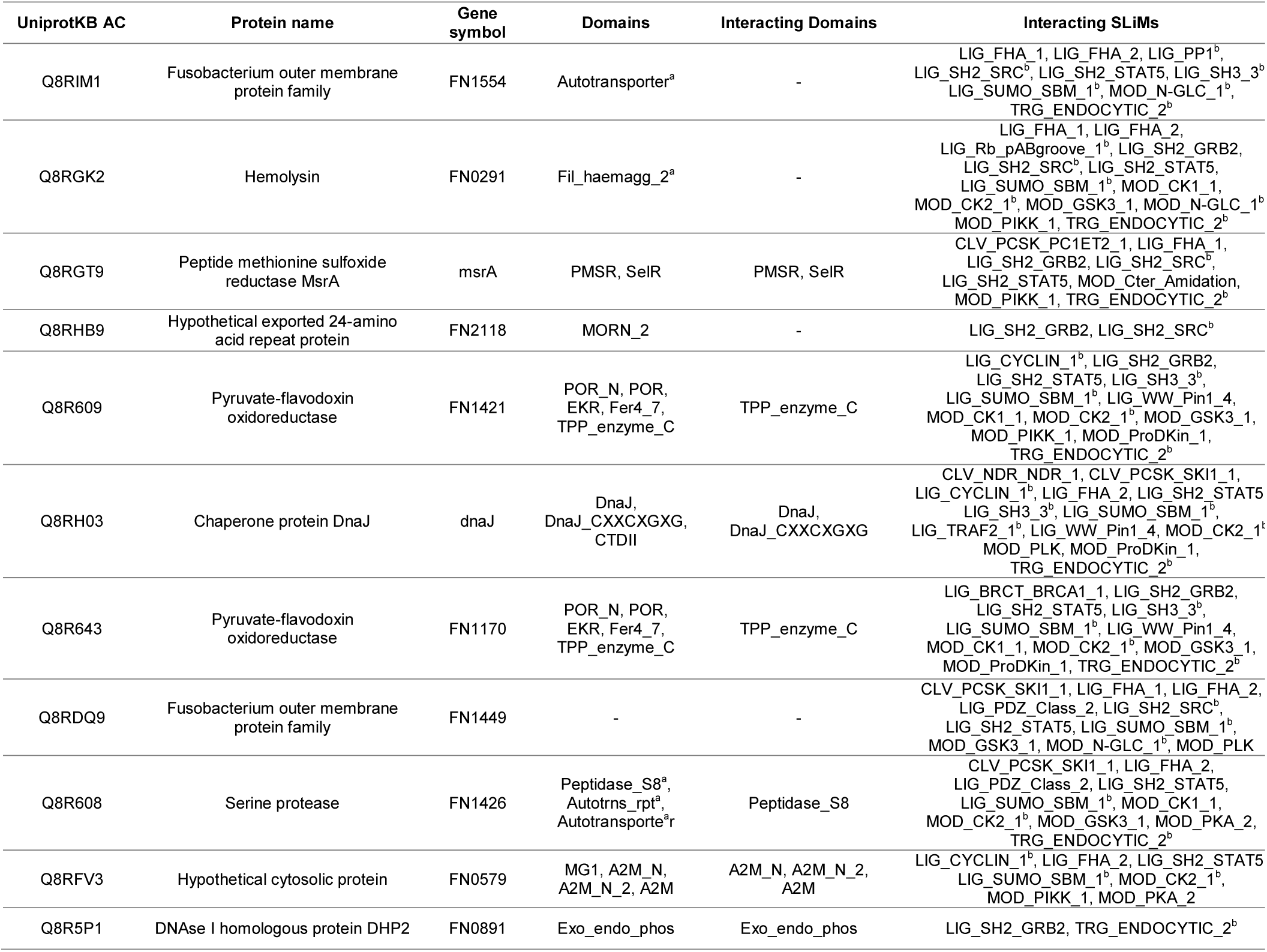
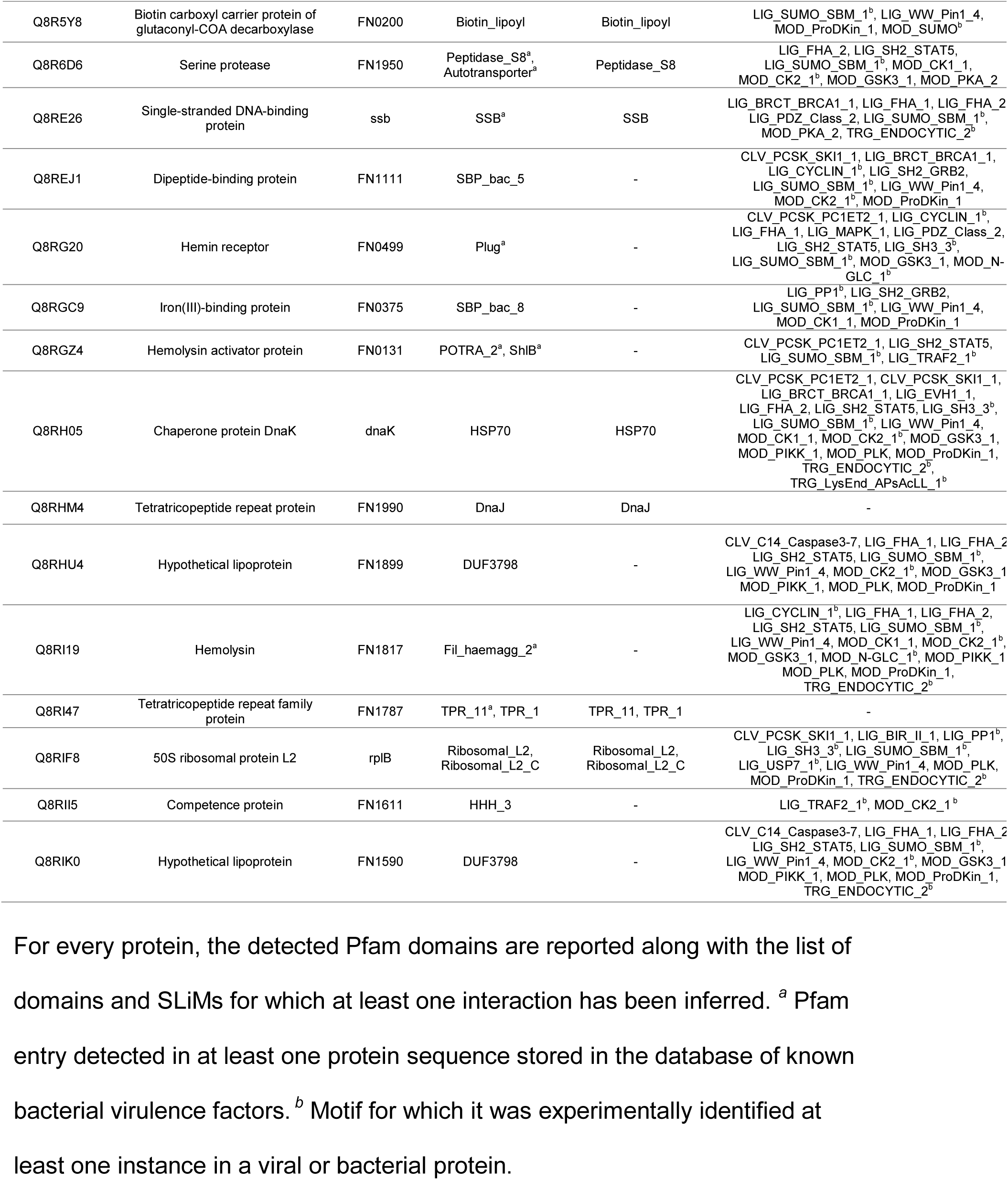
List of the main candidate virulence proteins in the *Fuso*Secretome.

### *F. nucleatum* and gut diseases from a network perspective

Among the 855 network modules detected in the human interactome, 38 are enriched in genes involved in at least one gut disease (*i.e.*, CRC and IBDs, see Methods). Interestingly, 27 of them (*i.e.*, 71%) are targeted by at least one *Fuso*Secretome protein, among which 3 contain a statistically significant fraction of inferred human interactors (Figure 5). Notably, Module 78, involved in immune response, is enriched in genes associated to inflammatory bowel diseases (IBDs) (28 proteins, 5.2-fold enrichment, P-value= 4.78×10^−4^) as well as in CD-specific (9 proteins, 13.4-fold enrichment, P-value = 2.52×10^−3^) and CRC-mutated (11 proteins, 4-fold enrichment, P-value = 1.46×10^−2^) genes. Moreover, it is enriched in genes whose expression correlates with *F. nucleatum* abundance in CRC patients (24 proteins, 3.7-fold enrichment, P-value = 3.35×10^−4^). This module is targeted by several main candidate virulence proteins, including a hemolysin (FN0291), an outer membrane protein (FN1554) and the MORN_2 domain containing protein (FN21118) (Figure 4A), which therefore, may play critical roles in these diseases. IBD genes are also enriched in Module 702 (5 proteins, 11-fold enrichment, P-value = 2.13×10^−2^), whose proteins participate in Jak-STAT signaling, whereas CD-specific genes are over-represented in Module 9 implicated in immunity (5 proteins, 28-fold enrichment, P-value = 2.52×10^−3^). Interestingly, Module 9 is specifically perturbed by Fap2 (FN1449) (Figure 4B), which is known to modulate the host immune response.

**Figure 5.**
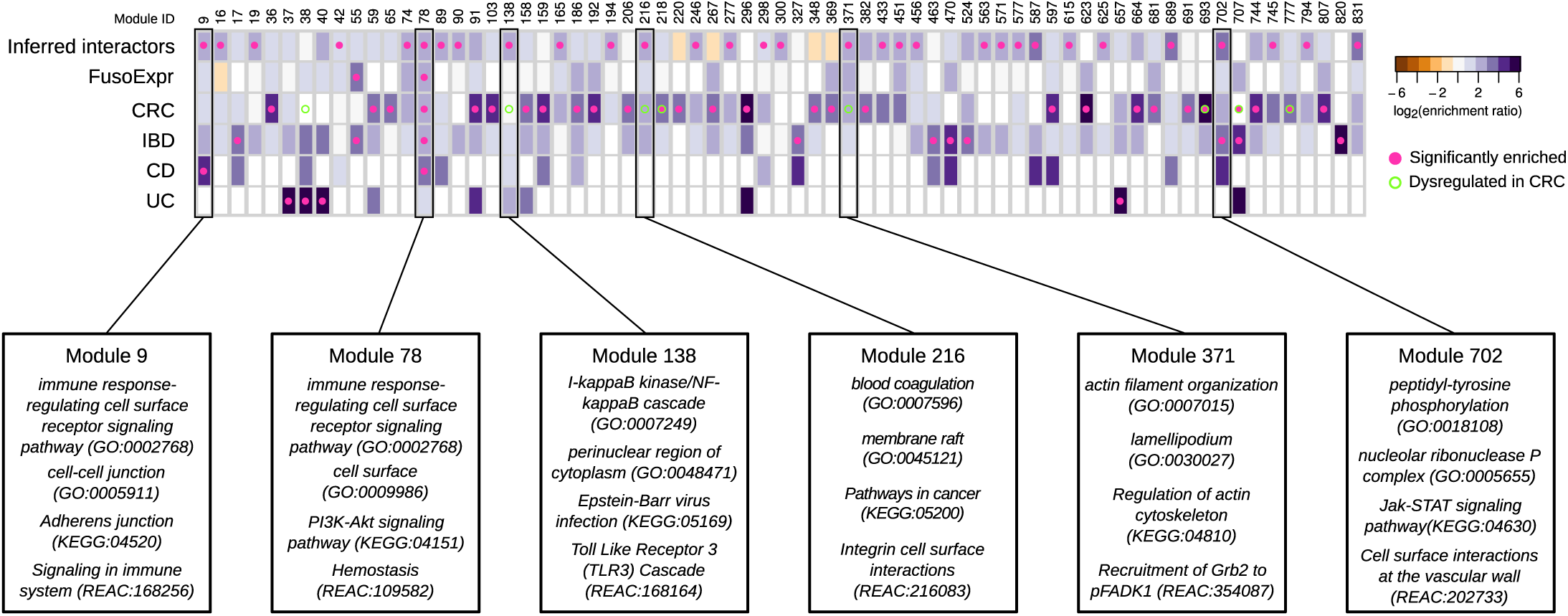
Enrichment of *Fuso*Secretome inferred human interactors and gut disease related proteins in network modules. Each column of the heatmap represents a module. The color of the cells corresponds to the log-transformed enrichment ratio. Pink circles indicate enriched sets. Modules showing a significant dysregulation in CRC progression are highlighted by an empty circle with green border. For the six modules showing an enrichment in inferred interactions and at least in one of gut disease related proteins, the most representative functions are reported. FusoExpr: genes whose expression correlates with *F. nucleatum* abundance in CRC patients; CRC: genes mutated in colorectal cancer samples; IBD: genes associated to inflammatory bowel disease; CD and UC: genes specifically associated to Crohn's disease and ulcerative colitis respectively.

Three other modules enriched in inferred human interactors show a significant dysregulation of the expression of their constituent proteins during CRC progression [66] and are implicated in infection-response pathways and cytoskeleton organization (Figure 5). In particular, two of these modules (Modules 138 and 216) show significant and specific up-regulation in Stage II, whereas the third (Module 371) is significantly up-regulated in normal and Stage II samples. Overall, these results indicate that *F. nucleatum* could contribute to the onset and progression of IBDs and CRC by perturbing some of the underlying network modules.

### Comparison with additional bacterial strains

We applied our computational approach on the recently released proteomes of 6 actively invading Fusobacteria strains isolated from biopsy tissues [8,65] (*i.e.*, 4 *F. nucleatum* subspecies and 2 *F. periodonticum* strains), and the proteome of *E. coli K-12* as a “control strain” (see Additional File 12: Supplementary Results, Table S12). We found that the secretomes of these 7 bacteria share common features (*i.e.,* disorder propensity, enriched domains, host-like domain and mimicry SLiM content) with the *Fuso*Secretome (Additional File 12: Table S13-S15 and Figure S2-S8). However, we observed a moderate overlap in terms of inferred interactors, enriched functions and preferentially targeted network modules (Additional File 12: Table S16-S18), and a modest concordance in term of network module perturbators (Additional File 12: Table S19).

The results of these analyses suggest that, on the one hand, actively invading Fusobacteria species share common mechanisms to interact with host cell, and, on the other hand, are consistent with the fact that *F. nucleatum* is an unusual heterogeneous species both at the genotypic and phenotypic level [8,65,67]. Finally, the commonalities between the *Fuso*Secretome and the *E. coli* K-12 secreted proteins are not surprising, since previous work showed that *E. coli* K-12 carries cryptic genes coding for virulence factors [68], whose expression is activated by mutations in the histone-like protein HU, which convert this established commensal strain to an invasive species in intestinal cells [69].

## Discussion

Over the years, it has been shown that *F. nucleatum* can adhere and invade human cells triggering a pro-inflammatory response. Nevertheless, the current knowledge on the molecular players underlying the *F. nucleatum* – human cross-talk is still limited.

For this reason, we carried out a computational study to identify *F. nucleatum* putative secreted factors (*Fuso*Secretome) that can interact with human proteins. The originality of our study is manifold compared to previous work. First, we used secretion prediction to identify potential *F. nucleatum* proteins that can be present at the microbe-host interface. Second, we exploited both domain-domain and domain-motif templates to infer interactions with human proteins. Earlier works, including one on *F. nucleatum*, chiefly applied homology-based methods for interaction inference with host proteins (*e.g.*, [70–73]). To our knowledge, domain-motif templates have been only exploited so far to infer or to resolve human-virus protein interaction networks [39,74]. Indeed, SLiM mimicry is widespread among viruses [21,75], but increasing evidence shows that it can be an effective subversion strategy in bacteria as well [22]. Third, we performed a network-based analysis on the human interactome to identify the main candidate *F. nucleatum* virulence proteins and the sub-networks they likely perturb.

Our approach relies on two prediction steps: *(i)* the definition of the *Fuso*Secretome based either on the presence of a signal peptide or several protein features such as disorder content, and *(ii)* the detection of host mimicry elements involved in the interaction with the host. It could be argued that the SecretomeP algorithm may incorrectly predict some proteins as secreted because of their high disorder content. For instance, a previous study considered as erroneous the secretion prediction of ribosomal proteins [76]. We assigned 20 ribosomal proteins to the *Fuso*Secretome. Although we cannot exclude a misprediction, ribosomal proteins can be secreted in some bacteria and be involved in host interaction [77,78]. Furthermore, increasing evidence shows that ribosomal proteins are moonlighting proteins with extra-ribosomal functions such as the *E. coli* ribosomal L2 protein that moonlights by affecting the activity of replication proteins [79]. Among the 337 inferred interactions between the 20 *Fuso*Secretome ribosomal and 183 human proteins, only a third of latter belong to ribosomal protein families. Interestingly, only 3 of the 41 human interactors inferred for *F. nucleatum* L2 are ribosomal proteins, and we identified the L2 protein as candidate virulence protein preferentially targeting Module 451. As this module is mainly involved in cell cycle and DNA repair, this result is consistent with the ability of L2 in *E. coli* to interfere with DNA processing factors [79] and further reinforces the confidence in the secretome prediction. Moreover, we have here underlined the value of the proposed approach: the interactome provides, on the one hand, the proper biological context to filter out potential false positive inferred interactions and, on the other, pinpoints candidate proteins that can be involved in the *F. nucleatum* – host interface.

Concerning the host mimicry elements, SLiM detection is notorious for over-prediction [54], given their relative short length and degeneracy (*i.e.*, few fixed amino acid positions). Our strategy to control for false positives was to consider only conserved SLiM occurrences in the *Fuso*Secretome protein regions predicted as disordered. Indeed, the vast majority of known functional SLiMs falls in unstructured regions [54,56] and shows higher levels of conservation compared to neighboring sequences. Conversely, we might also have missed some “true” mimicry instances in the *Fuso*Secretome by using too stringent parameters for domains and SLiMs identification and our interaction inferences may well be incomplete due to the limited number of available interaction templates. However, their functional significance fortifies our confidence in the predictive approach. Indeed, the *Fuso*Secretome shares similar features with known virulence proteins highlighting its pathogenic potential. In addition, interactors are implicated in established biological processes and cellular districts of the host-pathogen interface and significantly overlap with known pathogen protein binders. Furthermore, more than 70% of interactors are expressed in either the saliva or intestinal tissues. This suggests that most of the inferred interactions can occur in known *F. nucleatum* niches in the human body. Finally, we found among the human interactors an over-representation of genes whose expression correlates with *F. nucleatum* in CRC patients [13] as well as in IBD-related genes [80], which are mainly involved in immune-and infection-response pathways.

Moreover, we gained a broader view of the cellular functions that can be perturbed by the *Fuso*Secretome by investigating the human interactome. Although our interactome contains some functional inherent biases typical of literature-based interaction networks [81] (see Additional file 12), it better covers the interactions space of human secreted proteins, which are not easy to investigate using large-scale interaction screening methods such as yeast-two hybrid [82].

In agreement with previous experimental observations of host cell networks targeted by distinct pathogens, *F. nucleatum* targets hubs and bottlenecks in the human interactome [30,33,57]. Interestingly, the *Fuso*Secretome tend to interact with multifunctional proteins. This can represent an effective strategy to interfere with distinct cellular pathways as the same time [83].

Among the network modules preferentially targeted by the *Fuso*Secretome, we identified, besides the well-established functions related to host – pathogen interactions, several modules involved in chromatin modification and transcription regulation (Modules 246, 451, 571 and 625), and localized in compartments such as perinuclear region of the cytoplasm (Modules 90, 138 and 615). Intriguingly, this is reminiscent of the fact that invading *F. nucleatum* strains localize in perinuclear district of colorectal adenocarcinoma cells [8], and that bacteria can tune host-cell response by interfering directly – or indirectly – with the chromatin organization and the regulation of gene expression [84].

We propose 26 *Fuso*Secretome candidate virulence proteins as major network perturbators. They are the predominant interactors of preferentially targeted modules. Among the candidates we identified the known virulence protein Fap2, which was recently shown to promote immune system evasion by interacting with the immunoreceptor TIGIT [19]. Interestingly, Fap2 interacts specifically with Module 9, which is involved in immune response, thus suggesting novel potential binders mediating Fap2 subversion.

A recent report found that abundance of *F. nucleatum* is associated with high microsatellite instability tumors and shorter survival [14]. Notably, three preferentially targeted network modules (*i.e.*, Modules 138, 216 and 371) show a significant up-regulation in a stage associated to high microsatellite instability during CRC progression (stage II) [66] [85] and poor prognosis [86] [87]. This suggests that these modules may be important for CRC progression and outcome, and that the inferred interactions targeting these modules can mediate the cross-talk between *F. nucleatum* and the host in this particular subtype of CRC.

Overall, our functional and network-based analysis shows that the proposed interactions can occur *in vivo* and be biologically relevant for the *F. nucleatum* – human host dialogue.

## Conclusions

Over the last years, many microbes have been identified as key players in chronic disease onset and progression. However, untangling these complex microbe-disease associations requires lot effort and time, especially in the case of emerging pathogens that are often difficult to manipulate genetically. By detecting the presence of host mimicry elements, we have inferred the protein interactions between the putative secretome of *F. nucleatum* and human proteins, and ultimately provided a list of candidate virulence proteins and their human interactors that can be experimentally exploited to test new hypotheses on the *F. nucleatum* – host cross-talk. Our computational strategy can be helpful in guiding and speeding-up wet lab research in microbe-host interactions.

## Methods

### Protein sequence data

The reference proteomes of *Fusobacterium nucleatum subsp. nucleatum* strain ATCC 25586 (Proteome ID: UP000002521) and *Homo sapiens* (Proteome ID: UP000005640) were downloaded from the UniProtKB proteomes portal [88] (April 2013). The protein sequences of known gram-negative bacteria virulence factors were taken from the Virulence Factors DataBase [49] (January 2014).

### Secretome prediction

We identified putative secreted proteins among the *F. nucleatum* proteins by applying two algorithms: SignalP 4.1 [44] that detects the presence of a signal peptide and SecretomeP 2.0 [45] that identifies non-classical secreted proteins (i.e. not triggered by a signal peptide) using a set of protein features such as amino acid composition and intrinsic disorder content.

### Disorder propensity

To evaluate the intrinsic disorder propensity of *F. nucleatum* proteins predicted as secreted, we used the stand-alone programs of the following algorithms: DISOPRED (version 2.0) [89], IUPred (both long and short predictions) [90] and DisEMBL (COILS and HOTLOOPS predictions, version 1.4) [91]. We compared the disorder propensity distribution of SignalP-predicted secreted proteins to non-secreted proteins using the Kolmogorov-Smirnov test (two-sided, alpha = 0.05).

### Detection of functional domains

We ran the pfamscan program [92] on *F.nucleatum*, *H. sapiens* and virulence factors protein sequences to detect the presence of Pfam domains [52] (release 26). We kept only Pfam-A matches with an E-value < 10^−5^.

### Identification of short linear motifs

We used the SLiMSearch 2.0 tool from the SLiMSuite [93] to identify occurrences of known short linear motifs from the ELM database [53] (downloaded in May 2013) in the *F. nucleatum* proteome. To select putative mimicry motifs, we applied two SLIMSearch context filters: *(i)* the motif must be in a disordered region (average motif disorder score > 0.2, calculated by IUPred) and *(ii)* must be conserved in at least one putative ortholog detected in a database of 694 proteomes of commensal/pathogen bacteria in Mammalia downloaded from UniprotKB (March 2014). Sequence alignments and conservation assessment were performed using the GABLAM program from the SLiMSuite using standard parameters [94].

### Protein interaction inference

We built an interaction network between *F. nucleatum* putative secretome and human proteins by using interaction templates from the 3did database [95], which stores 6290 high-resolution three-dimensional templates for domain-domain interactions, and the iELM resource [96,97] that lists 578 high-confidence motif-mediated interfaces between 191 ELM motifs and 402 human proteins. Both datasets were downloaded in August 2013. The domain-based interaction inference works as follow: given a pair of known interacting domains *A* and *B*, if domain *A* is detected in the *F. nucleatum* protein *a* and domain *B* in the human protein *b*, then an interaction between *a* and *b* is inferred. Analogously, for the SLiM-mediated interaction inference: for a given known ELM motif *m* interacting with the domain *C* in the human protein *c*, if the motif *m* occurs in the *F. nucleatum* protein *a*, then *a* is inferred to interact with *c*.

### Human proteins targeted by bacteria and viruses

We gathered a list of 3,428 human proteins that were experimentally identified as interaction partners of three bacterial pathogen proteins (*Bacillus anthracis*, *Francisella tularensis*, and *Yersinia pestis*) in a large-scale yeast two-hybrid screen [30]. We downloaded interaction data with viruses for 4,897 human proteins from the VirHostNet database [98].

### Human expression data

RNA-seq expression data for 20,345 protein coding genes in normal colorectal, salivary gland and small intestine (i.e., jejunum and ileum) tissues was downloaded from the Human Protein Atlas (version 13), a compendium of gene and protein expression profiles in 32 tissues [99]. We considered as expressed those protein-coding genes with a FPKM > 1, that is 13,640 for colorectal, 13,742 for salivary gland and 13,220 for small intestine.

### Functional enrichment analyses

We have compiled several gut-related disease gene sets gathering data from the literature and public repositories. Patient secretome profiling (2,566 proteins) for tumor colorectal tissue samples were taken from [100]. We retrieve 152 colorectal cancer genes from the Network of Cancer Genes database (version 4.0, [101]). The list of human genes whose expression correlates with *F. nucleatum* abundance in colorectal cancer patients [13] was kindly provided by Aleksandar Kostic (Broad Institute, USA). The compendium of 163 loci associated with inflammatory bowel diseases was taken from a large meta-analysis of Crohn's disease and ulcerative colitis genome-wide association studies [80]. The enrichment of these gut-related disease gene sets among inferred interactors was tested using a one-sided Fisher's exact test.

We assessed the over-representation of cellular functions by performing a enrichment analysis on the list of inferred human interactors using the g:Profiler webserver [102] (version: r1488_e83_eg30, build date: December 2015). We analyzed the following annotations: Biological Process and Cellular Component from the Gene Ontology [103]; biological pathways from KEGG [59] and Reactome [60]. Functional categories containing less than 5 and more than 500 genes were discarded.

We used two different reference backgrounds for these statistical analyses. The first background consists of the protein-coding genes in the human genome (*i.e.*, 20'254 genes, UniprotKB, February 2013), whereas the second includes 11'284 protein-coding genes for which we could infer an interaction based on the available domain-domain and motif-domain interaction templates. In both cases, P-values were corrected for multiple testing with the Benjamini–Hochberg procedure applying a significance threshold equal to 0.025.

### Human interactome building, network module detection and annotation

We use the human interactome that we assembled and used in [62,66]. Briefly, protein interaction data were gathered from several databases (*e.g.*, BioGRID, InnateDB, Intact, MatrixDB, MINT, Reactome) through the PSICQUIC query interface [104] and from large-scale interaction mapping experiments (e.g., [105]). We kept only likely direct (*i.e.,* binary) interactions according to the experimental detection method [106] and mapped protein identifiers to UniprotKB IDs. Given the redundancy among SwissProt and TrEMBL entries, protein sequences were clustered using the CD-HIT algorithm [107]. SwissProt/TrEMBL pairs at 95% identity were considered as the same protein: interactions of TrEMBL protein were assigned to the SwissProt protein. As a result, we obtained a human binary interactome containing 74,388 interactions between 12,865 proteins (February 2013).

We detected 855 network modules detected using the Overlapping Cluster Generator algorithm [63]. Modules were functionally annotated by assessing the enrichment of Gene Ontology (GO) biological process and cellular component terms [103], and cellular pathways from KEGG [59] and Reactome [60]. Enrichment P-values were computed using the R package gProfileR [102] and corrected for multiple testing with the Benjamini–Hochberg procedure (significance threshold = 0.025) and annotated proteins in the human interactome were used as statistical background. Similarly, the over-representation of inferred human interactors and gut disease gene sets in network modules of the human interactome was assessed using a one-sided Fisher's exact test followed by Benjamini–Hochberg multiple testing correction (significance threshold = 0.025).

### Network module perturbation Z-score

We devised a score to quantify the contribution of *F. nucleatum* secreted proteins to the perturbation of a network module through their inferred interactions. We defined the perturbation Z-score for each *F. nucleatum* protein *f* interacting with at least one protein in module *m* as follows:

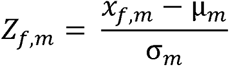

Where *x_f,m_* is the number of inferred interactions of the protein *f* with module *m*, *Z_f,m_* is the perturbation Z-score of the protein *f* in the module *m*, μ*_m_* and σ*_m_* are the mean of the inferred interaction values and their standard deviation in the module *m*, respectively.

### Network modules significantly dysregulated during CRC progression

The 77 network modules showing a significant dysregulation during CRC progression were taken from our previous work [66], in which we devised a computational method that combines quantitative proteomic profiling of TCGA CRC samples, protein interaction network and statistical analysis to identify significantly dysregulated cellular functions during cancer progression.

### Availability of data and material

All data generated or analyzed on the ATCC 25586 strain are included in this published article (and its supplementary information files). All other data are available from the corresponding author on reasonable request.

## Competing interests

The authors declare that they have no competing interests.

### Funding

The project leading to this publication has received funding from Excellence Initiative of Aix-Marseille University - A*MIDEX, a French “Investissements d'Avenir” programme and was partially supported by the French ‘Plan Cancer 2009–2013’ program (Systems Biology call, A12171AS). The funding organizations had no role in the design of the study and collection, analysis, interpretation of data, and in writing the manuscript.

### Author contributions

AZ conceived the study, designed and performed the experiments, analyzed the data and wrote the manuscript. LS performed the experiments and analyzed the data. SB performed the experiments. CB designed the experiments, analyzed the data and wrote the manuscript. All authors read and approved the final manuscript.

## Acknowledgements

The authors would like to thank the members of the TAGC laboratory for fruitful discussion, Anaïs Baudot (I2M, CNRS, France) for critically reading the first draft of the manuscript, Aleksandar Kostic (Broad Institute, USA) for kindly providing the list of human genes whose expression correlates with *F. nucleatum* abundance in colorectal cancer patients, and Henrik Nielsen (DTU Bioinformatics, Denmark) for assistance in running SecretomeP predictions. AZ is grateful to Coralie, Olivia and Claire for their constant support.

## Consent for publication

Not applicable.

## Ethics approval and consent to participate

This study is based on publically available datasets only. Thus, no ethical approval is needed/applicable, nor is consent from any participants.

